# The iron-binding siderophore enterobactin is required for the response of multi-drug resistant *Klebsiella pneumoniae* to zinc limitation

**DOI:** 10.64898/2026.09.26.754270

**Authors:** Jeffrey R. Singer, D. Annie Doyle, Dillon E. Kunkle, Owen S. Burroughs, Rachel J Service, Walter J. Chazin, Eric P. Skaar

## Abstract

To persist during infection *Klebsiella pneumoniae* must overcome nutrient iron and zinc limitation imposed by the host immune system through a process called nutritional immunity. Secreted small molecule siderophores are a major virulence determinant of *Klebsiella pneumoniae* pathogenesis and are presumed to overcome nutritional immunity by binding iron for bacterial acquisition. In this work, we set out to identify how a multi-drug resistant *K. pneumoniae* grows in zinc limited environments. Using unbiased transcriptomics, proteomics, and an arrayed transposon screen, we identified that synthesis and uptake of the siderophore enterobactin is required to allow for growth in low zinc conditions. Iron-specific chelators did not replicate this phenotype and addition of supplemental iron through heme in growth media could not complement severe growth defects of enterobactin mutant *K. pneumoniae* experiencing zinc limitation. Finally, zinc starvation induced enterobactin production independent of the canonical zinc uptake regulator (Zur) transcription factor suggesting an unidentified regulatory mechanism by which Gram-negative pathogens may respond to zinc stress. Together, these studies expand the role of enterobactin beyond iron regulation and highlight a previously unreported link between iron and zinc homeostasis in *Klebsiella pneumoniae*.

**IMPORTANCE:** Enterobactin is the archetypal model for understanding siderophore-mediated iron acquisition in Gram-negative bacteria and has the highest affinity for iron of any known molecule. Here, we identify an essential biological role for enterobactin acquisition in response to nutrient zinc limitation in an ST258 strain of *Klebsiella pneumoniae* and identify the enterobactin biosynthetic gene cluster as a Zur-independent locus of regulation to bacterial zinc stress. These findings expand the role of enterobactin in nutritional immunity and identify a new regulatory mechanism for *Klebsiella pneumoniae* zinc homeostasis.

## INTRODUCTION

*Klebsiella pneumoniae* (*Kpn*) is a Gram-negative opportunistic pathogen of the Enterobacteriaceae family recently labeled by the World Health Organization as the highest priority pathogen for antimicrobial research and development due to rising multi-drug resistance seen globally (MDR)(1). The majority of antibiotics in clinical practice currently target a short list of highly conserved processes: bacterial cell wall biogenesis, nucleic acid synthesis, or protein synthesis(2). Another conserved biological function common to all organisms is the acquisition of nutrient metal. Approximately 30% of all proteins in nature require metals to carry out their biological function, therefore all bacterial pathogens must acquire metals from their host during infection (3). The “trojan horse” antibiotic cefiderocol exploits this requirement by linking a cephalosporin moiety to a siderophore backbone to enhance bacterial uptake of the cell wall biogenesis inhibitor(4). Still, a better understanding of pathogen metal acquisition strategies may inform novel approaches for antimicrobial development.

Selective pressure for microbial evolution of high affinity metal acquisition is imposed by a repertoire of host proteins expressed in immune cells that specifically chelate iron (Fe) and other metals through a process coined nutritional immunity (5). Calprotectin (CP) is a dominant nutritional immunity effector protein with high affinity binding to zinc (Zn), Fe, copper (Cu), nickel (Ni) and manganese (Mn)(6). CP is highly abundant at sites of infection as it is estimated to account for 45% of the cytoplasmic protein in a neutrophil (7).

One strategy that *Enterobacteriaceae* have evolved to obtain nutrient metals from a metal-limited environment is through the enzymatic production and uptake of small molecule metal-binding compounds called siderophores. Most siderophores have even higher affinity for metal than CP with K_a_ measured as high as 10^52^ (6, 8). Siderophore generation and import are known to be virulence determinants in multiple pathogens including *Kpn* (9–11). Interestingly, *Kpn* has undergone a bifurcation in evolutionary trajectories with some “hypervirulent” strains acquiring multiple accessory plasmid-encoded siderophores while “classical” strains have less redundant Fe acquisition systems and instead are permissive to antibiotic resistance plasmids (12). Among classical *Kpn* strains, the siderophore enterobactin is ubiquitously encoded, while yersiniabactin has more variable representation (13).

While numerous studies have described a role for Fe acquisition in *Kpn* pathogenesis, less is known about *Kpn* adaptations to Zn limitation. *Kpn* encodes two functionally redundant high affinity ATP-Binding Cassette (ABC) transporters through operons *znuCBA* and *zniCBA* which contribute to Zn homeostasis and pathogenesis in a murine model of pneumonia (14). Interestingly, proteomic studies of a different strain of *Kpn* grown in Zn-limited vs. Zn-replete media did not identify increased abundance of either transporter (15). In *E. coli*, *A. baumannii*, and other Gram-negative organisms, *znuCBA* expression is controlled by de-repression of the Zinc Uptake Regulator (Zur) (16, 17). Although *Kpn* encodes for a Zur homolog, it is not known if the same regulatory mechanisms are evolutionarily conserved across species.

In this work, we aimed to identify factors essential for *Kpn* to overcome nutrient Zn limitation using unbiased and orthogonal genome-wide approaches of RNA-sequencing (RNA-seq), whole cell proteomics, and an arrayed-transposon library screen. RNA-seq and proteomics coupled to this analysis in a *zur::Tn* mutant create a powerful dataset to not only identify genes important for Zn limitation, but also identify if they are likely regulated by Zur or another transcription factor in *Kpn*. This work highlights one such example where we identify a novel and surprising role for the Fe-binding siderophore enterobactin whose uptake and production are induced through Zn limitation, required for growth in low Zn conditions, but whose regulation is independent of Zur.

## RESULTS

### Identification of genes that are required for *Kpn* growth in response to Zn limitation

To study the response of MDR *Kpn* to nutrient Zn limitation, the ST258 strain KPNIH1 (18) was grown in vehicle alone or increasing concentrations of the Zn chelator N,N,N’N’-Tetrakis(2-pyridylmethl) ethylenediamine (TPEN). A concentration of 50µM was identified that inhibits growth but does not completely restrict growth (Data not shown). We hypothesized that the most attractive genes for study that were important for *Kpn* to contend with nutrient Zn stress would be transcriptionally upregulated and increased in protein abundance following exposure to a Zn chelator. However, differential expression alone does not necessarily imply functional importance and therefore an arrayed-transposon library screen was also completed to identify genes whose disruption impacted *Kpn* growth in low Zn conditions.

Following 15 minute treatment with TPEN or vehicle, RNA was extracted from bacterial cells and subjected to RNA-Seq. Compared with vehicle treated cells, treatment with TPEN resulted in 106 genes that were greater than 2-fold increased with adjusted p-value< 0.5 (Figure 1A). In accordance with microarray studies in *E. coli* treated with TPEN (19, 20), the most upregulated genes were known to encode putative Zn importers (*znuA*, *zniA*, *zinT*), ribosomal subunits involved in Zn homeostasis (*rpmJ2* and *rpmE2*), and Fe acquisition systems (*entCEBA*, *hmuSTUV*, *sitABC*). Since transcriptional changes do not always correlate well with proteome abundance (20) and post-transcriptional regulation is an important aspect of pathogen virulence (21), untargeted proteomics using whole-cell lysates was conducted to further identify which of the 106 upregulated genes were increased in abundance at the protein level.

**Figure 1:**
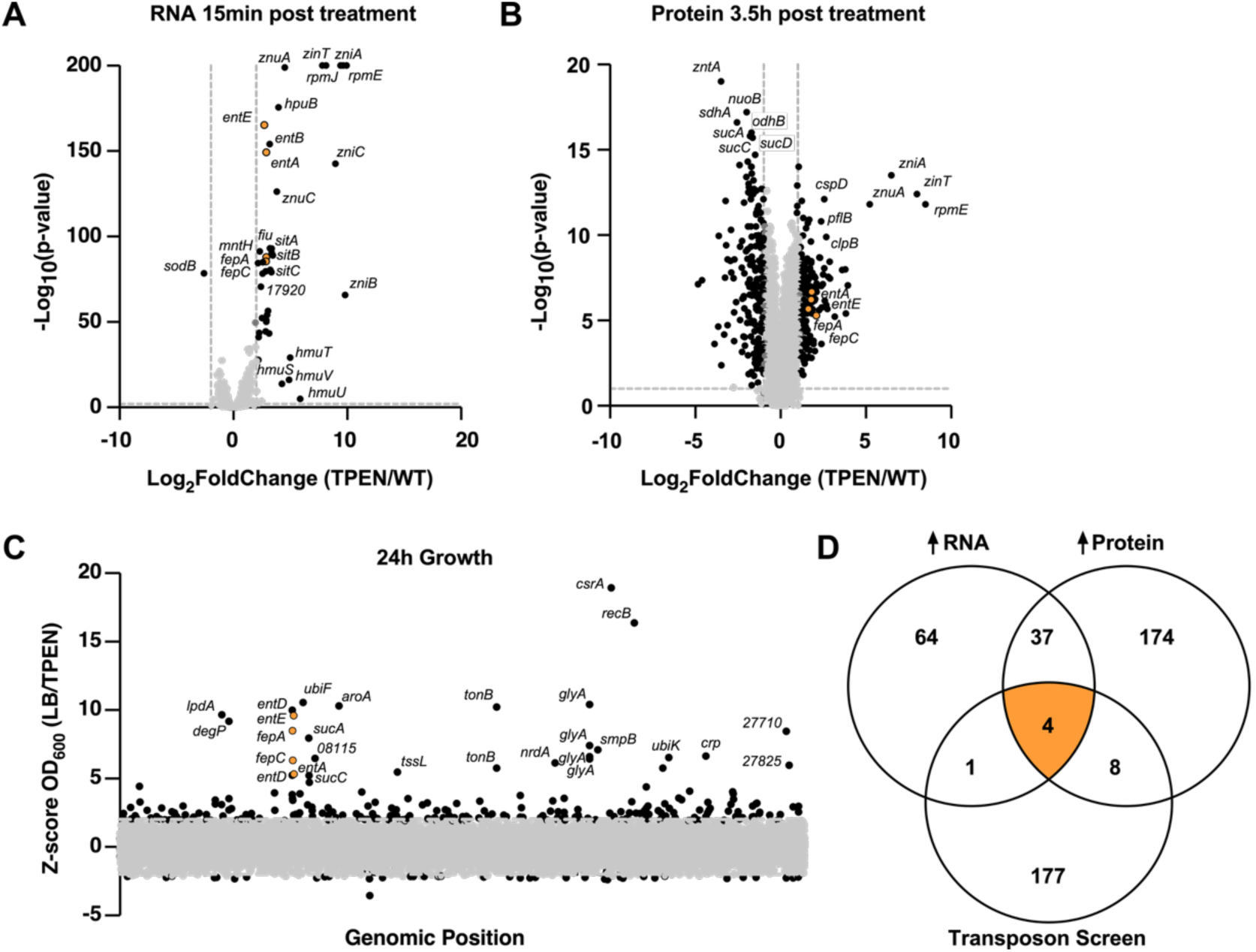
Identification of genes that are required for *Kpn* growth in response to Zn limitation. (**A**) Differential gene expression of RNA transcripts and **(B**) Differential abundance of proteins in whole cell proteomics from 4 biological replicates of *Kpn* treated with vs. vehicle (100% ethanol) vs. 50 µM TPEN. Each dot represents a single gene within *Kpn* genome. Vertical dotted lines denote 2-fold change in expression/abundance respectively. Horizonal line denote adjusted p-value of 0.05. (**C**) Arrayed-transposon library screen of *Kpn*. Each dot represents the Z-score of the ratio of OD_600_ measured after 24 hours of growth in an individual transposon mutant treated with vehicle or 50µM TPEN. Some genes contain multiple mutants. X-axis displays genomic position and. Horizontal dotted lines indicate cutoff of □2 SD from the mean. Genes were labeled if they were greater than 5 SD from mean OD_600_ ratio. (**D**) Venn diagram depicting genes whose expression/abundance were increased greater than 2-fold with adjusted p-value <.05 after treatment of 50µM TPEN and/or Z-score of OD_600_ ratio >2 SD from the mean. 4 genes (*entA, entE, fepA, fepC*) identified across all experiments are highlighted in orange in each figure.

Vehicle or TPEN treated bacteria were grown to mid-log phase, 3 vs. 3.5 hours respectively, and OD-matched pellets were lysed, protein isolated, and subjected to LC-MS/MS for proteomics analysis. Using a 2-fold increase and adjusted p-value of 0.05 as cutoffs, 223 proteins were identified as increased in abundance following TPEN treatment. The proteins most increased in abundance were also genes with the highest transcriptional upregulation which provides internal validation for the dataset (Figure 1B). These proteins include Zn importers ZnuA, ZniA, ZinT and ribosomal subunit RpmE2. However, only 18.3% of enriched peptides following TPEN treatment were also upregulated in the RNA-seq dataset further indicating a large role for post-transcriptional adaptation to nutrient Zn stress.

To determine which of the 41 genes upregulated transcriptionally and at the protein level were required for growth in Zn limitation, a KPNIH1 arrayed-transposon library (18) was screened for growth after 24h incubation in the same 50µM concentration of TPEN. This library was generated to contain a total of 12,000 mutants each with a randomly inserted mini-Tn5 derivative transposon carrying chloramphenicol resistance. Prior characterization of the library identified approximately 85% of predicted protein-coding genes having transposon mutagenesis with an average of 2.5x coverage per gene. Since multiple mutants with transposon insertions disrupting the same gene were represented across the library on different plates, an additional layer of reproducibility was built into the screen. The arrayed-transposon library screen identified 205 bacterial mutant hits representing transposon insertions across 190 genes indicating the majority of hits had only 1 mutant represented (Figure 1C). As not every mutant has been whole-genome sequenced in this library to confirm its identity, we considered candidate genes for further study if any of the represented mutants showed severe growth inhibition in TPEN.

Comparison across all genomic datasets identified only 4 genes upregulated transcriptionally and with increased protein abundance after TPEN treatment whose disruption led to severe growth restriction in TPEN treated medium: *entE, entA*, *fepA*, and *fepC* (Figure 1D). *entD*, a gene in the same operon as *fepA* was transcriptionally upregulated and also identified as a hit in the transposon screen, but protein abundance did not meet significance threshold. Surprisingly, these genes are all located at the same genetic locus (Figure 2A) and encode for the synthesis and uptake of the siderophore enterobactin. Perhaps even more remarkably, but in accordance with Maunders et al., disruption of all known Zn importers through transposon mutagenesis was insufficient to cause growth delay in Zn-limited media (14).

**Figure 2:**
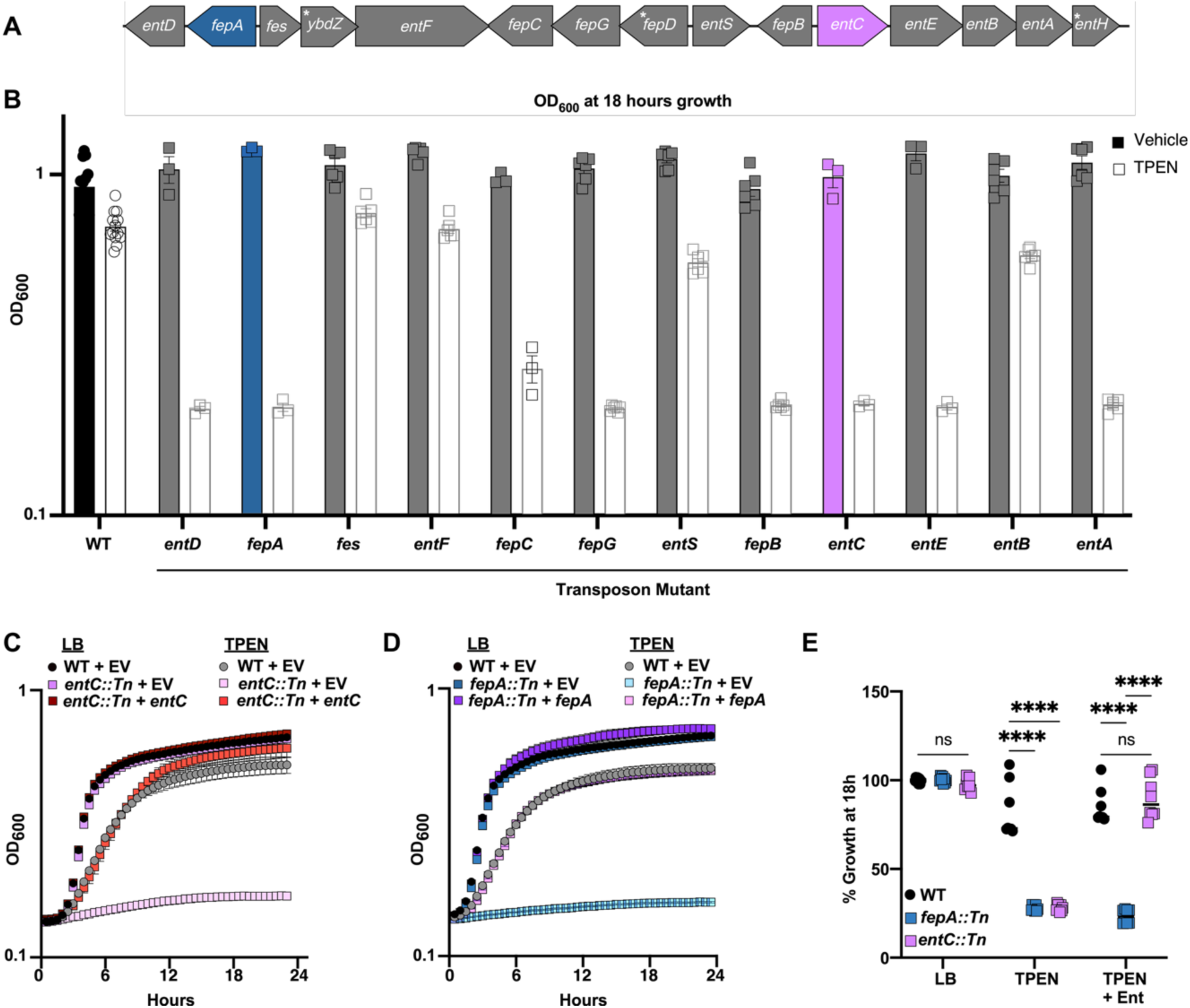
Enterobactin uptake and production are required for growth after TPEN treatment. (**A**) Cartoon representation of *Kpn* genomic locus that encodes the biosynthetic gene cluster for enterobactin production and uptake. Genes with * denote that they are not represented in arrayed-transposon library generated by Ramage et al. (**B**) WT *Kpn* or indicated transposon insertion mutant were grown in the presence of vehicle (solid bars) or 50µM TPEN (empty bars) and OD_600_ was monitored every 30 minutes. OD_600_ at 18 hours is indicated. Data pooled from 3-14 biological replicates. Error bars show □ SEM. WT, (**C**) *entC::Tn,* or (**D**) *fepA::Tn Kpn* harboring empty vector (EV) control plasmid or plasmid encoding indicated gene were cultures in LB medium alone or 50µM TPEN and OD_600_ was monitored every 30 minutes for 24 hours. (**E**) WT, *fepA::Tn*, or *entC::Tn Kpn* were cultured in LB medium alone, 50µM TPEN, or 50µM TPEN and 15µM apo-enterobactin and OD_600_ was monitored every 30 minutes for 24 hours. At 18 hours, OD_600_ values were normalized to WT *Kpn* untreated. Data pooled from 8 biological replicates in technical duplicate across 2 separate experiments. Two-way ANOVA with Tukey’s multiple comparisons test. **** p <.0001

Enterobactin is a tricyclic catecholate siderophore produced by Enterobacteriaceae family organisms like *Salmonella*, *Escherichia*, *Yersinia*, and *Klebsiella* and has the highest binding affinity for Fe^3+^ of any known natural or synthetic chemical (22). Since its discovery in 1970, numerous elegant studies predominantly in *E. coli* have identified the structural and regulatory mechanisms by which bacteria utilize this small molecule to obtain Fe for a myriad of cellular processes (22). While *E.* coli-derived enterobactin has also been identified to have a role in reduction of copper leading to toxic stress (23, 24) and *Rothia*-produced enterobactin is hypothesized to bind Zn and magnesium (25), little else is known about its role in bacterial physiology outside of iron acquisition and no direct role shown in response to Zn stress.

### Enterobactin is required for *Kpn* growth after treatment with TPEN

To validate results from the arrayed-transposon library screen, individual mutants were isolated from wells in the arrayed library and assessed for growth in Zn limitation with bacterial growth curves. Confirmatory severe growth inhibition was observed in mutants for *entD*, *fepA*, *fepC*, *entE*, and *entA* treated with TPEN. Additionally, *fepG, fepB*, and *entC* mutants also showed similar growth restriction despite not being identified as “hits” on the transposon library screen (Figure 2B). The production of enterobactin occurs through a multi-enzymatic reaction of a nonribosomal peptide synthetase complex from the amino acid precursor chorismate (22). Following conversion from chorismate into 2,3-dihydroxybezoic acid (DHB) via EntC, EntB, and EntA, an amide linkage between DHB and L-serine is catalyzed by entE, entD, and entF. Three DHB-serine molecules then cyclize to form the ringed siderophore structure where Fe^3+^ can coordinate between the deprotonated catechol groups (22). Once excreted into the periplasm through EntF, the multi-drug exporter TolC facilitates cellular export where enterobactin can bind Fe^3+^. The holo-enterobactin is then recognized by the TonB-dependent outer membrane protein FepA. Back in the periplasm, enterobactin binds FepB before the ABC transporter FepDGC transports enterobactin across the inner membrane. Finally, *fes* encodes a hydrolase to cleave the trilactone backbone and liberate Fe for intracellular use (22).

While all components of enterobactin uptake are required to overcome growth restriction in TPEN, disruptions of *entF* and *entB* via transposon mutagenesis did not lead to severe growth restriction. Similarly, transposon mutants with *fes* or *entS* disrupted also grew similar to WT treated with TPEN. While prior work identifying clinical isolates with 347bp deletion in *entS* identified it is not critical for catecholate siderophore export (26) the other results are not easily explained via polar effects or within a model that precursors of enterobactin can overcome its loss since EntB has previously been deleted in *Kpn* to disrupt enterobactin production (27). Taken together, these data suggest while complex, the production and uptake of enterobactin are required for growth following treatment with TPEN. To simplify experimental design and interpretation, subsequent studies utilized WT and two transposon mutant strains with no growth inhibition in rich media, but severe growth restriction in 50µM TPEN if the initial step of either biosynthesis (*entC:;Tn*) or import (*fepA::Tn*) of enterobactin were disrupted (Figure 2C). Growth could be complemented in either mutant strain with a full-length copy of the disrupted gene and its endogenous promoter delivered in trans (Figure 2D). Further, *entC::Tn* mutant growth in Zn limitation was able to be complemented with exogenously provided enterobactin, while *fepA::Tn* mutant remained growth restricted in the presence of TPEN indicating that enterobactin import and not intracellular biosynthesis is required to withstand Zn limitation (Figure 2E).

### Enterobactin production and import are regulated by Zn limitation independent of Zur

TPEN is a commonly used reagent to apply intracellular Zn stress through chelation, but it is not entirely specific to Zn with known affinity for Fe, Mn, and Cu (28). Indeed, given the multiple copies of high affinity Zn importers in *Kpn* we wondered if TPEN’s growth restriction was exerted by Zn limitation or through another trace transition metal like Fe. Transcriptional and proteomic data suggested global upregulation of Fe acquisition systems following TPEN treatment and the role of enterobactin is primarily understood to be specific to Fe import (19, 22). Therefore, intracellular metal concentration was quantified following treatment with chelators. Mid-log WT or *fepA::Tn* mutant bacteria were treated with vehicle, 50µM TPEN, or 350µM of a more specific Fe chelator 2’2-dipyridyl (DIP) for 1 hour before ICP-MS. Reduction in intracellular Zn was observed only in bacteria treated with TPEN, but both TPEN and DIP led to decrease in intracellular Fe (Figure 3A). However, despite similar or larger reduction in intracellular Fe following DIP treatment compared to TPEN, *fepA::Tn* and *entC::Tn* mutants did not experience severe growth restriction with 350µM DIP treatment (Figure 3B). These results indicated that Fe restriction alone is unlikely to account for the lack of fitness of enterobactin production or uptake mutants following treatment with TPEN.

**Figure 3:**
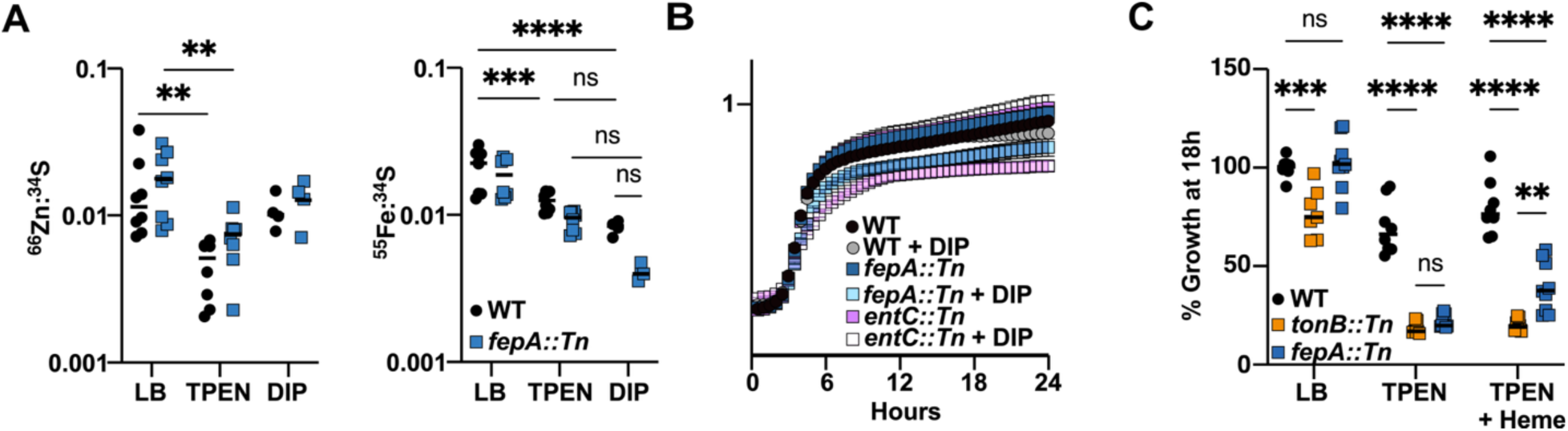
Fe limitation does not explain severe growth restriction of *Kpn* after TPEN treatment. (A) Total cellular zinc (Zn) and iron (Fe) concentrations in WT or *fepA::Tn Kpn* strains cultured in LB medium for 3 hours prior to being left untreated (LB) or treated with either 50µM Zn chelator TPEN or 350µM Fe chelator Dipyridyl for an additional one hour. Each point represents a biological replicate in technical triplicate. Two way ANOVA with Tukey’s multiple comparisons test ** p < .01, *** p < .001, **** p < .0001. (B) WT, *fepA::Tn*, or *entC::Tn Kpn* were cultured in LB medium alone or 350µM Fe chelator Dipyridyl and OD_600_ was monitored every 30 minutes for 24 hours. (**C**) WT, *tonB::tn*, or *fepA::Tn Kpn* were cultured in LB medium alone, 50µM TPEN, or 50µM TPEN and 10µM Heme and OD_600_ was monitored every 30 minutes for 24 hours. At 18 hours, OD_600_ values were normalized to WT *Kpn* untreated. Data pooled from 8 biological replicates in technical duplicate across 2 separate experiments. Two-way ANOVA with Tukey’s multiple comparisons test. ** p <.005, *** p <.001, **** p <.0001.

If severe growth limitation exerted by TPEN was due to Fe chelation, we hypothesized that supplemental Fe should recover growth defects in TPEN-treated bacteria. However, because TPEN binds Fe cations directly, interpretation is challenging because growth recovery after Fe treatment could alternatively be explained if chelator saturation prevented TPEN binding of another metal exerting a biologic effect. To explore whether Fe could recover growth following TPEN treatment, Fe inaccessible to TPEN was supplemented through the form of heme. Heme is an Fe-binding small molecule porphyrin ring utilized across all kingdoms of life including bacterial pathogens (29). The TonB*-*dependent Hmu uptake system was increased in transcriptomic and proteomic datasets following TPEN treatment of *Kpn*, so we reasoned exogenous heme could be readily utilized by *Kpn* for Fe-dependent cellular processes. Indeed, TPEN treated *fepA::Tn* mutant *Kpn* was able partially restore growth with 10µM exogenous heme compared to *tonB::Tn* mutant *Kpn* unable to import both heme and enterobactin (Figure 3C). These results further cast doubt on a model where the role of enterobactin was solely dependent on Fe acquisition.

To investigate whether enterobactin played a role in Zn limitation independent of Fe chelation, more precise tools were required than industrial small-molecule chelators like TPEN and Dipyridyl. Extensive prior work has validated the use of recombinant CP and site-specific mutant CP with altered metal-binding properties to more accurately control nutrient metal limitation in microbial pathogenesis (30–33). CP forms a heterodimer of S100A8 and S100A9 proteins with two canonical metal-binding sites at the protein-protein interface (34, 35). One site is comprised of 6 histidine side chains and binds Zn, Cu, Mn, Ni, and Fe with high affinity while the other site forms a tetrahedral His_3_Asp coordination with only Zn and Cu to high affinity (30). After being treated with 500µg/mL CP, WT *Kpn* showed moderate growth inhibition which prevented growth completely in *entC::Tn* or *fepA::Tn* mutants (Figure 4A). Deletion of both His_6_ and His_3_Asp (CP^ΔS1/S2^) sites preventing metal binding capabilities of CP restored *Kpn* growth to levels observed culture bacteria in buffer alone. Interestingly, H103/104/H105N (CP^H3N^) mutant which is able to bind Zn and Cu with high affinity, but not Mn, Fe, or Ni phenocopied CP albeit at a higher dose of 1000µg/mL suggesting Zn and/or Cu chelation alone is sufficient for severe growth restriction in the absence of enterobactin production or uptake. Severe growth restriction imposed by CP^H3N^ in enterobactin transposon mutants could be restored to inhibition of WT *Kpn* with 15µM enterobactin supplementation in mutants able to import enterobactin, *entC::Tn*, but not in *fepA::Tn* (Figure 4B). Taken together, these data demonstrate enterobactin uptake is required for growth in nutrient Zn limitation.

**Figure 4:**
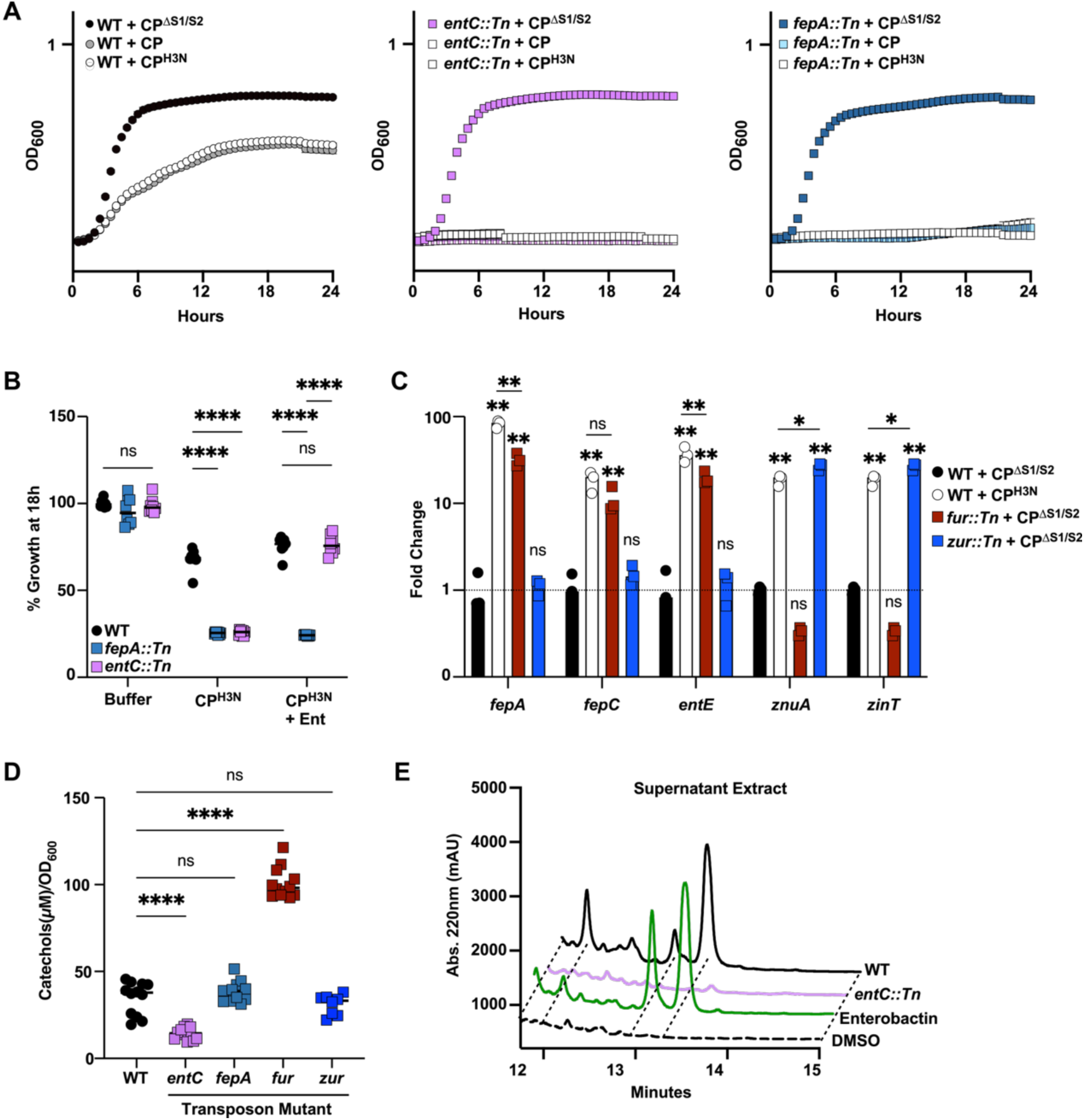
Enterobactin uptake and production are induced in Zn limitation and required for growth independent of Zur. (**A**) WT, *entC::Tn,* or *fepA::Tn Kpn* were cultured in LB and calprotectin (CP) buffer with addition of 500µg/mL WT CP, 1000µg/mL inactive CP (CP^ΔS1/S2^), or 1000µg/mL Zn-specific CP (CP^H3N^) and OD_600_ was monitored every 30 minutes for 24 hours. Data pooled from 8 biological replicates in technical duplicate across 2 separate experiments. (**B**) WT, *fepA::Tn*, and *entC::Tn* strains of *Kpn* were cultured in LB and CP buffer alone, 1000µg/mL Zn-specific CP (CP^H3N^), or 1000µg/mL Zn-specific CP (CP^H3N^) with addition of 15µM apo-enterobactin and OD_600_ was monitored every 30 minutes for 24 hours. At 18 hours, OD_600_ values were normalized to WT *Kpn* untreated in CP buffer alone. Two-way ANOVA with Tukey’s multiple comparisons test. **** p <.0001. (**C**) qPCR of indicated transcript abundance in WT, *fur::Tn*, or *zur::Tn Kpn* after 1 hour treatment with CP^ΔS1/S2^ or CP^H3N^ where indicated. Fold change was calculated using ΔΔCT method normalized to WT *Kpn* treated with inactive CP^ΔS1/S2^. Two-way ANOVA with Tukey’s multiple comparisons test. Asterix above condition is compared to WT + CP^ΔS1/S2^. * p < .01, ** p <.0001. (**D**) OD_600_ of overnight culture was measured and supernatant collected from indicated *Kpn* strains to assay for catechol siderophore using arnow’s test. One-way ANOVA with Tukey’s multiple comparisons test. **** p < .0001. (**E)** Supernatant from WT (black) or *entC::Tn* (pink) cultures grown in minimal media were methanol extracted and analyzed by HPLC. 30µM enterobactin (green) and equivalent DMSO (dashed line) in minimal media were extracted as control.

The major regulator of intracellular Zn limitation in *Kpn* is the transcription factor Zur which upon Zn binding attaches to “Zur box” DNA sites upstream of target genes to repress their transcription(17). When intracellular Zn is limited, Zur is derepressed allowing transcription of genes important for bacteria to contend with low nutrient Zn stress. The major regulator for enterobactin biosynthesis and uptake is a related transcription factor, Ferric Uptake Regulator (Fur). Three “Fur boxes” occupy promoters in the enterobactin biosynthetic gene cluster (BGC) upstream of divergent transcriptional start sites of *fepA/fes*, *fepD/entS*, and *entC/fepB*. No Zur boxes were bioinformatically predicted in the enterobactin BGC using motifs identified in *Kpn* MGH 78578 by RegPrecise (36) however, Zur boxes were correctly predicted for known Zur-target genes *znuCBA*, *zniCBA*, and *zinT* validating the approach.

To evaluate empirically if Zn limitation altered enterobactin BGC expression WT *Kpn* was treated with either CP^ΔS1/S2^ or CP^H3N^ for 1 hour prior to transcriptional analysis using quantitative PCR using ΔΔCT method normalizing to *recA* for reference gene (37). In addition to experimental groups, transcriptional changes were also investigated in *fur:Tn* and *zur:Tn* transposon mutants treated with CP^ΔS1/S2^ to add context to expression changes observed (Figure 4C). During Zn limitation imposed by CP^H3N^, genes across the enterobactin BGC were upregulated 20-80 fold which was equivalent or greater than maximal Fur derepression for all genes assessed. Conversely, no change in enterobactin BGC expression was observed in the *zur::Tn* mutant indicating transcriptional control of enterobactin in low Zn conditions is independent of Zur. To confirm Zn limitation in the assay, *znuA* and *zinT* transcript abundances were measured in Zn limitation at nearly 20-fold increase above CP^ΔS1/S2^ treated *Kpn* which was comparable, but significantly lower than transcript abundances measured with maximal Zur derepression in the *zur::Tn* mutant. While Zur is a transcriptional regulator, it is conceivable that a transcriptionally-independent action of Zur could lead to increased enterobactin production. Therefore, enterobactin production was measured in supernatant indirectly using Arnow’s assay (38) and directly following methanol extraction and high-performance liquid chromatography to confirm that while enterobactin production is under regulation from Fur and *entC:Tn* mutants do not secrete the siderophore, *zur::Tn* mutants did not produce increased enterobactin (Figure 4D & E). Collectively, these results suggest Zn limitation and not Fe limitation induces gene expression changes to produce and import enterobactin independent of Zur and establish a causal role for enterobactin in maintenance of *Kpn* Zn homeostasis.

Despite several studies identifying mechanisms of Zn homeostasis in *Kpn*, the Zur regulon has not previously been elucidated. Therefore, transcriptomic and proteomic studies in *zur::Tn* mutant were compared with WT *Kpn* to identify high confidence operons likely to be under control of the Zur regulon. 182 genes were upregulated compared to WT *Kpn* greater than 2-fold with adjusted p <.05. Amongst the most differentially expressed genes were *KPNIH1_05910*, *KPNIH1_05915*, and *KPNIH1_05920* encoding for two 50S ribosomal peptides Rpme2 and Rpmj2 recently identified in the *Neisseria gonorrhea* Zur regulon (39) and a predicted Gcn5-related N-acetyltransferase family protein (Figure 5A). Other highly Zur repressed genes include *zniCBA*, *znuCBA*, *zinT*, and *KPNIH1_17920* which encodes a predicted protein belonging to the COG0523 candidate Zn metallochaperone family (40, 41). While the definition of a regulon is inherently restricted to transcription factor binding and transcriptional regulation and not protein abundance, we hypothesized that transcriptionally upregulated and differentially abundant proteins between WT and *zur::Tn Kpn* would represent high confidence genes under direct repression by Zur. Unbiased proteomics identified 43 gene products increased with derepression of Zur, 11 of which were also upregulated transcriptionally (Figure 5B & C). While genes in the enterobactin BGC were uniformly upregulated in *zur::Tn Kpn* with adjusted p values <0.5, only the gene encoding the Fe-liberating enterobactin esterase *fes* was increased greater than 2-fold. Further, no differences were detected at the protein level across bacterial genotypes for the enterobactin BGC.

**Figure 5:**
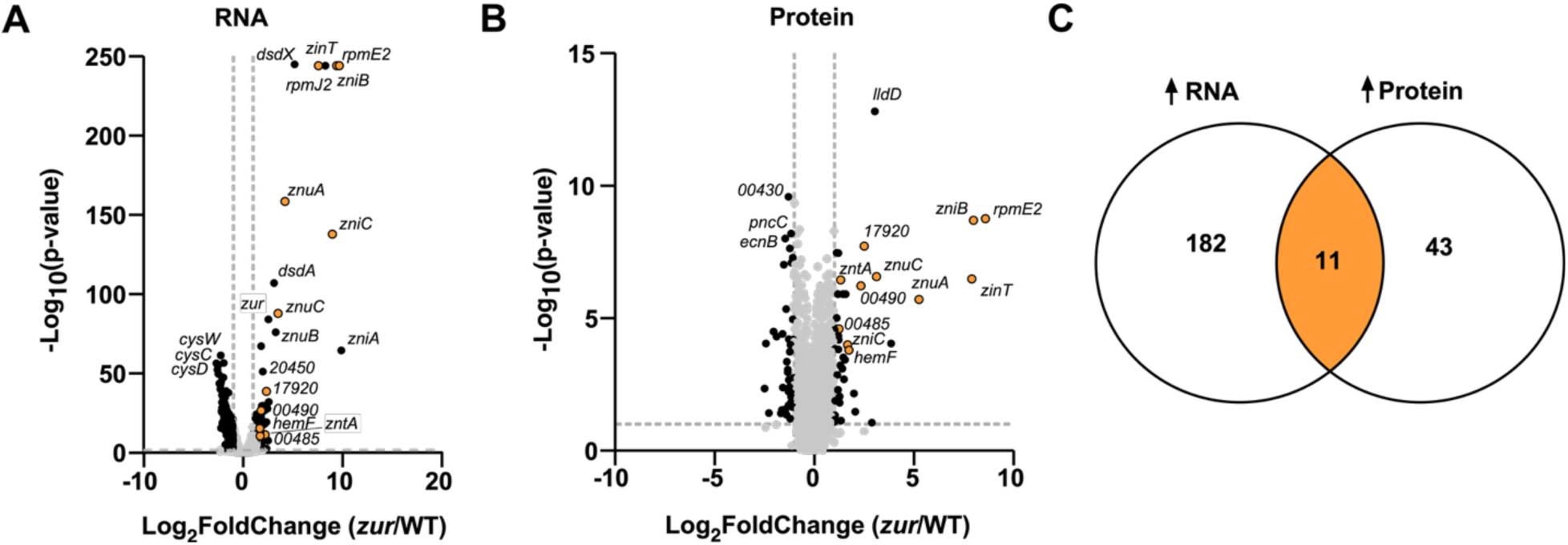
Transcriptional and proteomic changes following Zur derepression. (**A**) Differential gene expression of RNA transcripts and **(B**) Differential abundance of proteins in whole cell proteomics from 4 biological replicates of WT vs. *zur::Tn Kpn* grown to mid-log. Each dot represents a single gene within *Kpn* genome. Vertical dotted lines denote 2-fold change in expression/abundance respectively. Horizonal lines denote adjusted p-value of 0.05. (**C**) Venn diagram depicting genes whose expression/abundance were increased greater than 2-fold with adjusted p-value <.05 in *zur::Tn* mutant compared to WT *Kpn*. 11 genes identified across both techniques are highlighted in orange in each panel.

## DISCUSSION

A major objective of the host immune system during a bacterial infection is to restrict pathogen nutrient acquisition and thereby prevent growth, a process called nutritional immunity(5). To acquire Fe in a host-limited environment, *Kpn* and other pathogens produce small-molecules called siderophores that selectively bind Fe and allow microbial-specific Fe import (22). Nutritional immunity encompasses metal limitation beyond Fe and recent studies in *Yersinia* and *Salmonella* suggest siderophores may also be important in responding to nutrient Zn limitation (32, 42, 43). The specific mechanisms that *Kpn* uses to overcome Zn limitation are not known since it expresses multiple high affinity Zn transporters and genetic deletion of both receptor complexes produces only modest growth defect in Zn limited media(14). Using orthogonal genome-wide approaches, we uncovered an unexpected requirement for import of the iron-binding siderophore enterobactin to overcome Zn limitation in a MDR ST258 strain of *Kpn*.

As the archetypal siderophore expressed in *E. coli* and other Gram-negative Enterobacteriaceae, enterobactin biosynthesis and uptake have been extensively studied in Gram-negative pathogens that express enterobactin including hypervirulent and classical strains of *Kpn* (22, 44–46). Even bacteria that do not synthesize enterobactin express receptors for uptake and xenosiderophore utilization like pathogens *Acinetobacter baumannii*(*47*) and *Pseudomonas aeruginosa*(*48*) or commensal *Bacteroides thetaiotaomicron*(*49*). However, to-date no prior investigations have identified a biological role of enterobactin in Zn homeostasis. Enterobactin isolated from *Rothia* was experimentally determined to bind Zn ∼2.5 fold less tightly than EDTA which has K_d_ ∼10^-16^(25, 50) but if this interaction is biologically relevant has not been evaluated.

Whether the role of enterobactin in overcoming Zn limitation is due to direct Zn binding and import or secondarily related to Fe starvation remains unknown. Experimental design with a chelator that binds both Zn and Fe with high affinity like TPEN prevents disentangling the possibly interconnected role of Fe and Zn at a molecular level. Similarly challenging is the use of minimal media because *entC::Tn* and *fepA::Tn* mutants grow robustly in Chelex-treated M9 media (data not shown). Treating bacterial strains with recombinant CP and its engineered mutants offers an experimental approach for future studies to evaluate the molecular mechanisms underlying how enterobactin can restore growth in Zn limitation. In this work, the approach was critical to show enterobactin transcriptional regulation due to Zn limitation, but independent of Zur. It is important to consider that while utilized as a Zn-specific reagent, CP^H3N^ can bind Cu. However, a role for Cu restriction by CP has not been previously appreciated for bacterial pathogens in rich media although it has been observed for *Candida albicans* in 50% serum (51). Additionally, TPEN treatment does not impact intracellular Cu concentrations by ICP (data not shown).

The studies described heavily utilize an arrayed-transposon library for screening and hit validation of siderophore biology (18). Such collections are not without caveats. Indeed, several transposon mutants were not originally identified as having restricted growth in TPEN until they were individually isolated and assayed perhaps owing to isolate contamination at some point in the original experiment. Additionally, genetic complementation was feasible solely replacing the single gene disrupted through transposon mutagenesis and did not require complementation with downstream genes suggesting genes disrupted by T30 transposon in operons may still produce transcript allowing faithful gene expression further downstream. Evidence of this phenomenon is found in RNA reads mapping to *zur* in the *zur::Tn* mutant following RNA-seq (Figure 5A). Validation and whole-genome sequencing is therefore critical for rigorous discovery utilizing this resource.

Future studies are needed to determine whether loss of enterobactin production or import impacts classical *Kpn* pathogenesis *in vivo*. Prior work in hypervirulent *Kpn* strains that express accessory siderophores aerobactin, yersiniabactin, and salmochelin found enterobactin production is dispensable in pulmonary or nasopharyngeal infection when other siderophores are being produced (10, 11, 46). However, without functional redundancy of siderophore-mediated metal uptake, enterobactin may play a more outsized role in pathogenesis of MDR *Kpn*. Additionally, whether pathogenic *E. coli* that also only express enterobactin shows a similar Zn-specific phenotype is also interesting to consider. During infection, Zn, Fe, and other nutrient metals are simultaneously sequestered by the host, so it is possible that in addition to metal-specific responses from Fur or Zur, a shared metal starvation response is required for pathogenesis.

## MATERIALS AND METHODS

### Bacterial strains and growth conditions

Bacterial strains used in this study are indicated in Table 1. Other than experiments for Arnow’s assay, bacteria were grown in liquid lysogeny broth (LB) or on LB mixed with 1.5% agar (LBA). Gentamicin was used at 10µg/mL for plasmid maintenance and screening. For siderophore measurement and extractions, minimal M9 media was prepared with 1x M9 salts, 0.4% glucose, 1% casamino acids in deionized H_2_O. 1g/L Chele× 100 resin was added for 16h prior to vacuum filtration and subsequent addition of sterile-filtered 0.1mM CaCl_2_ and 1mM MgSO_4_. All bacterial growth in static or shaking incubators at 37°C and 180rpm.

**Table 1:** Bacterial Strains used in this study.

| Strain | Genotype | Gene ID; transposon library item number; location | Reference |
| --- | --- | --- | --- |
| MKP103 | WT | WT KPNIH1 strain used for transposon library generation | (17) |
| MKP103 | *entD::Tn* | KPNIH_07170; KP03313; tnkp1_lr150110p20q102 | (17) |
| MKP103 | *fepA::Tn* | KPNIH_07175; KP03317; tnkp1_lr150131p24q177 | (17) |
| MKP103 | *fes::Tn* | KPNIH_07180; KP03319; tnkp1_lr150214p09q172 | (17) |
| MKP103 | *entF::Tn* | KPNIH_07190; KP03324; tnkp1_lr150110p19q154 | (17) |
| MKP103 | *fepC::Tn* | KPNIH_07195; KP03322; tnkp1_lr150214p33q190 | (17) |
| MKP103 | *fepG::Tn* | KPNIH_07200; KP03323; tnkp1_lr150214p07q192 | (17) |
| MKP103 | *entS::Tn* | KPNIH_07210; KP03324; tnkp1_lr150110p19q154 | (17) |
| MKP103 | *fepB::Tn* | KPNIH_07215; KP03327; tnkp1_lr150214p34q179 | (17) |
| MKP103 | *entC::Tn* | KPNIH_07220; KP03329; tnkp1_lr150214p30q109 | (17) |
| MKP103 | *entE::Tn* | KPNIH_07225; KP03331; tnkp1_lr150131p12q192 | (17) |
| MKP103 | *entB::Tn* | KPNIH_07230; KP03334; tnkp1_lr150117p11q156 | (17) |
| MKP103 | *entA::Tn* | KPNIH_07235; KP03335; tnkp1_lr150124p06q130 | (17) |
| MKP103 | *tonB::Tn* | KPNIH_15670; KP06458; tnkp1_lr150214p04q105 | (17) |
| MKP103 | *fur::Tn* | KPNIH_07710; KP03523 ; tnkp1_lr150117p05q170 | (17) |
| MKP103 | *zur::Tn* | KPNIH_01460; KP00685; tnkp1_lr150131p19q116 | (17) |

### RNA-sequencing and data analysis

Overnight cultures of WT or *zur::Tn* strains were sub-cultured at 1:1000 dilution and grown for 3.5h to mid-log growth in LB medium. Ethanol was added to *zur::Tn* mutant and either ethanol or 50µM N,N,N’N’-Tetrakis(2-pyridylmethl) ethylenediamine (TPEN) were added to WT *Kpn* with 4 biological replicates and allowed to grow for an additional 15 minutes before being pelleted, resuspended in Trizol, and frozen at xya80C until RNA extraction. RNA was extracted using chloroform and subsequent RNEasy column isolation (Qiagen) with on-column DNaseI digest (Qiagen) per manufacturer’s instructions. RNA quality determined with Tapestation analysis through VANTAGE and purified RNA sent to SeqCenter LLC for library preparation and sequencing. Raw paired-end reads were trimmed and quality filtered using fastp (version 0.24.0) and aligned to KPNIH1 *K. pneumoniae* genome using HISAT2 (version 2.2.1) with spliced alignment disabled. Read counts were obtained using featureCounts function of Subread (version 2.0.8) and differential expression using DESeq2 (version 1.40.2) with default parameters was used to analyze either WT vs. TPEN or Wt vs. *zur::Tn* using R Version 4.4.1.

### Proteomic sample preparation

Overnight WT or *zur::Tn Kpn* cultures were grown in 4 biologic replicates and sub-cultured at 1:1000 dilution into 10mL LB with 50µM TPEN or equal volume of ethanol and grown to OD_600_ of 1. *zur::Tn* was not grown in LB and ethanol. 3.5mL of OD-matched culture was washed twice in PBS prior to pellet being frozen at −80C. Bacterial pellets were thawed in cold lysis buffer (150mM NaCl, 20mM Tris-HCl Ph7.5) with addition of cOmplete™ protease inhibitor (Millipore Sigma) and transferred to lysing matrix B tubes (MP Biomedicals) and bead beat at 6.0 M/S for 45 seconds. Equal volume lysis buffer with 10% SDS (5% SDS total) were added with additional bead beating 6.0M/S for 45 seconds. After centrifugation, lysate was removed and protein quantitated with BCA assay. For each sample, 10 µg of protein was brought to 40 µL in 5% SDS and reduced with 10 mM TCEP at 55 °C for 15 min. Proteins were then alkylated with 20 mM iodoacetamide for 30 min at room temperature in the dark. Samples were acidified to 2.5% phosphoric acid and diluted with six volumes of S-Trap binding buffer. They were then digested with 1 µg trypsin (1:10 enzyme:protein) for 1 h at 47 °C on micro S-Trap columns (ProtiFi) following the manufacturer’s protocol (52). Eluted peptides were dried, desalted on C18 StageTips(53) and resuspended in 0.1% formic acid. Peptide concentrations were measured by absorbance at 205 nm (NanoDrop). Samples were adjusted to 50 ng/µL in 0.1% formic acid containing 0.015% n-dodecyl-β-D-maltoside (DDM).

### LC-MS/MS analysis

Peptides (50 ng) were separated on a PepSep column (25 cm × 75 µm, 1.5 µm particles; Bruker) held at 50 °C, using a nanoElute 2 nanoflow HPLC system (Bruker) at 600 nL/min. Solvent A was 0.1% formic acid in water and solvent B was 0.1% formic acid in acetonitrile. The gradient ran from 3% to 28% B over 30 min and then to 37% B over 5 min, followed by a wash at 85% B. Eluting peptides were analyzed on a timsTOF HT mass spectrometer (Bruker) using data-independent acquisition with parallel accumulation–serial fragmentation (diaPASEF) (54). MS1 spectra were acquired from m/z 100 to 1700, with a 50 ms trapped ion mobility ramp covering 1/K0 0.75–1.30 V·s/cm². The diaPASEF scheme used 36 variable-width isolation windows (three per TIMS ramp across 12 ramps) spanning m/z 350–1250. Collision energy was ramped linearly with ion mobility from 20 eV at 1/K0 0.6 to 59 eV at 1/K0 1.6.

### Proteomic data analysis

Proteomic data were analyzed in Spectronaut v19.6 (Biognosys) (55) using the directDIA workflow. Each analysis searched a protein database from KPNIH1 circular genome and plasmids which contained 5,398 entries. Precursor and protein identifications were filtered to a 1% q-value. Protein groups were quantified using both MS1 and MS2 signal. Differential abundance between conditions was assessed with unpaired t-tests at the protein-group level, and p-values were adjusted by the Benjamini–Hochberg method.

### Arrayed Transposon Library Screen

Individual plates of MKP103 transposon mutants were thawed and disposable pin replicator (Scinomix) used to transfer colonies onto 96 well plate filled with 100µL LB/Agar and let grow at 37°C for approximately 24 hours. The following day, individual colonies were inoculated into 500µL of LB in deep 96-well plates and shaken at 180 rpm at 37°C overnight with Breath-Easy® sealing membrane. After overnight growth, each well was sub-cultured at 1:50 dilution for 1h in LB before diluted again 1:100 in 100µL LB with 100% ethanol or 50µM TPEN in round bottom 96 well plate. OD_600_ was measured across each plate at 24 hours using the same plate reader. OD_600_ values were imported to a dataframe in using R Version 4.4.1. using “wellr” package. Ratio of OD_600_ in vehicle (ethanol) divided by OD_600_ in TPEN-treated mutants was calculated and Z-score of this ratio was determined across all mutants tested.

### Bacterial growth curves

Overnight cultures of bacterial strains were started from single colonies on LBA into LB media (with antibiotics if appropriate). The following morning, culture was back-diluted 1:50 and grown at 37°C shaking at 180rpm for 1 hr. Back-diluted culture were then inoculated 1:100 into 96-well microtiter plates of indicated medium in technical duplicates in 100µl final volume. Breath-Easy® sealing membrane was added to prevent evaporation and edge-effects on plate. Plates were cultured for 24h at 37°C with continuous shaking and OD_600_ measurements recorded every 30 minutes. CP and all variants were prepared as previously reported (31, 56). WT CP or mutant protein was diluted to 2.5x final concentration (500µg/mL or 1000µg/mL respectively) in filter-sterilized CP buffer (20mM Tris-HCl pH 7.5, 100mM NaCl, 3mM CaCl_2_, and freshly added 5mM 2-mercaptoethanol). 40µl CP Buffer per well was added to 59µl LB and mixed prior to addition to 1µl back-diluted bacterial culture. For enterobactin add-back experiments, initial concentration of 25µM apo-enterobactin (Sigma) was prepared in LB media for final concentration of 15µM with addition of CP Buffer.

For Heme growth assays, 10mM stock hemin was prepared in 0.1M NaOH before being diluted 1:1000 to a final concentration of 10µM for bacterial growth.

For add-back experiments, OD_600_ was examined at 18 hours into growth curve in all bacterial wells and normalized to WT *Kpn* in vehicle treated sample as 100% maximum growth.

### Cloning and plasmid construction

All plasmids and primers used in this study are indicated in Table 2 and Table 3 respectively. Plasmids were constructed using directional cloning with T4 ligase or HiFi Assembly per manufacturer’s protocol. Prior to use, Plasmid constructs were sequence verified with Plasmidsaurus Whole Plasmid Sequencing (Oxford Nanopore, R10.4.1). Complementation vector backbone was constructed by PCR amplification of backbone of pACYC184 (ATCC) excluding chloramphenicol resistance gene and inserting PCR-amplified gentamicin resistance from pUC18T-mini-Tn7T-Gm using HiFi DNA Assembly mastermix (NEB) with indicated primers.

**Table 2:**
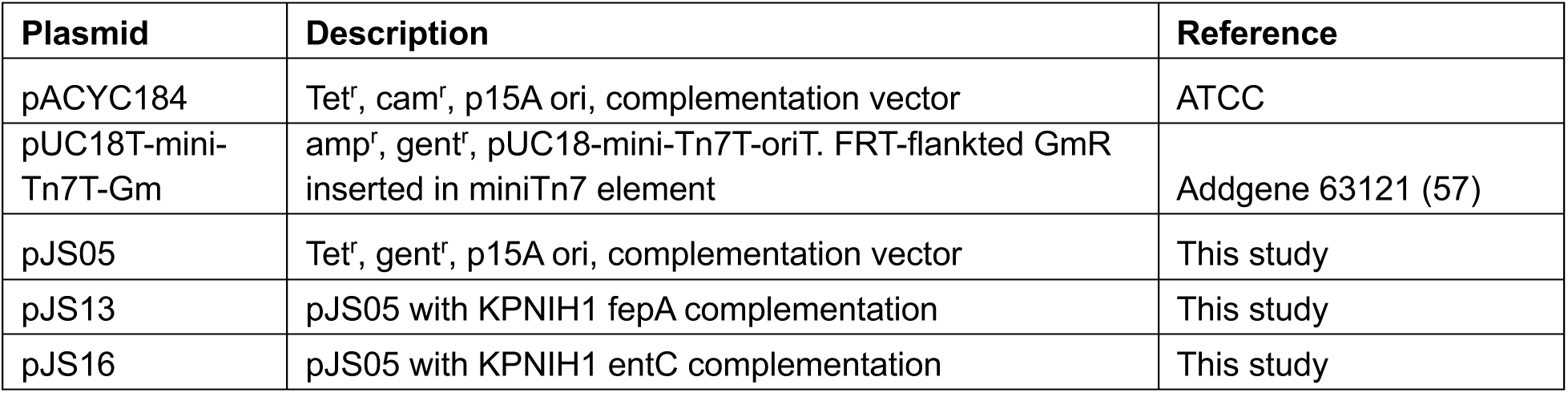
Plasmids used in this study.

| Plasmid | Description | Reference |
| --- | --- | --- |
| pACYC184 | Tet<sup>r</sup>, cam<sup>r</sup>, p15A ori, complementation vector | ATCC |
| pUC18T-mini-Tn7T-Gm | amp<sup>r</sup>, gent<sup>r</sup>, pUC18-mini-Tn7T-oriT. FRT-flanked GmR inserted in miniTn7 element | Addgene 63121 (57) |
| pJS05 | Tet<sup>r</sup>, gent<sup>r</sup>, p15A ori, complementation vector | This study |
| pJS13 | pJS05 with KPNIH1 *fepA* complementation | This study |
| pJS16 | pJS05 with KPNIH1 *entC* complementation | This study |

**Table 3:**
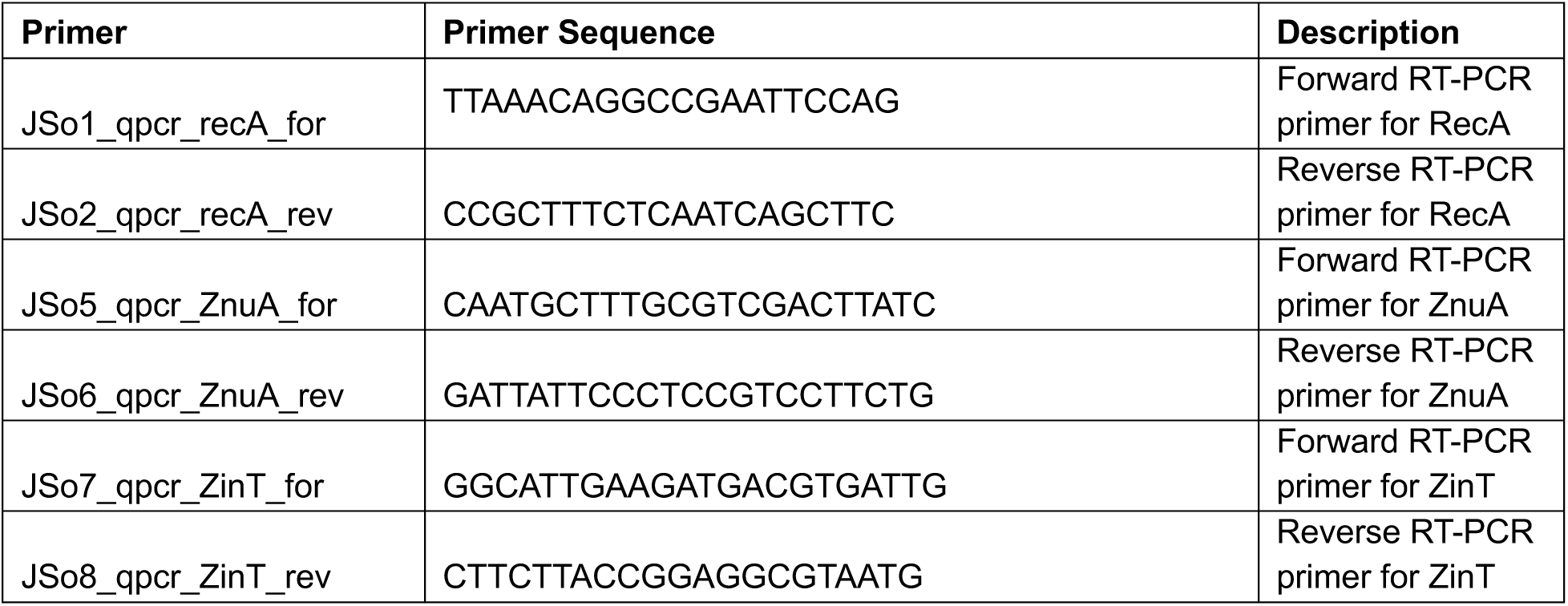
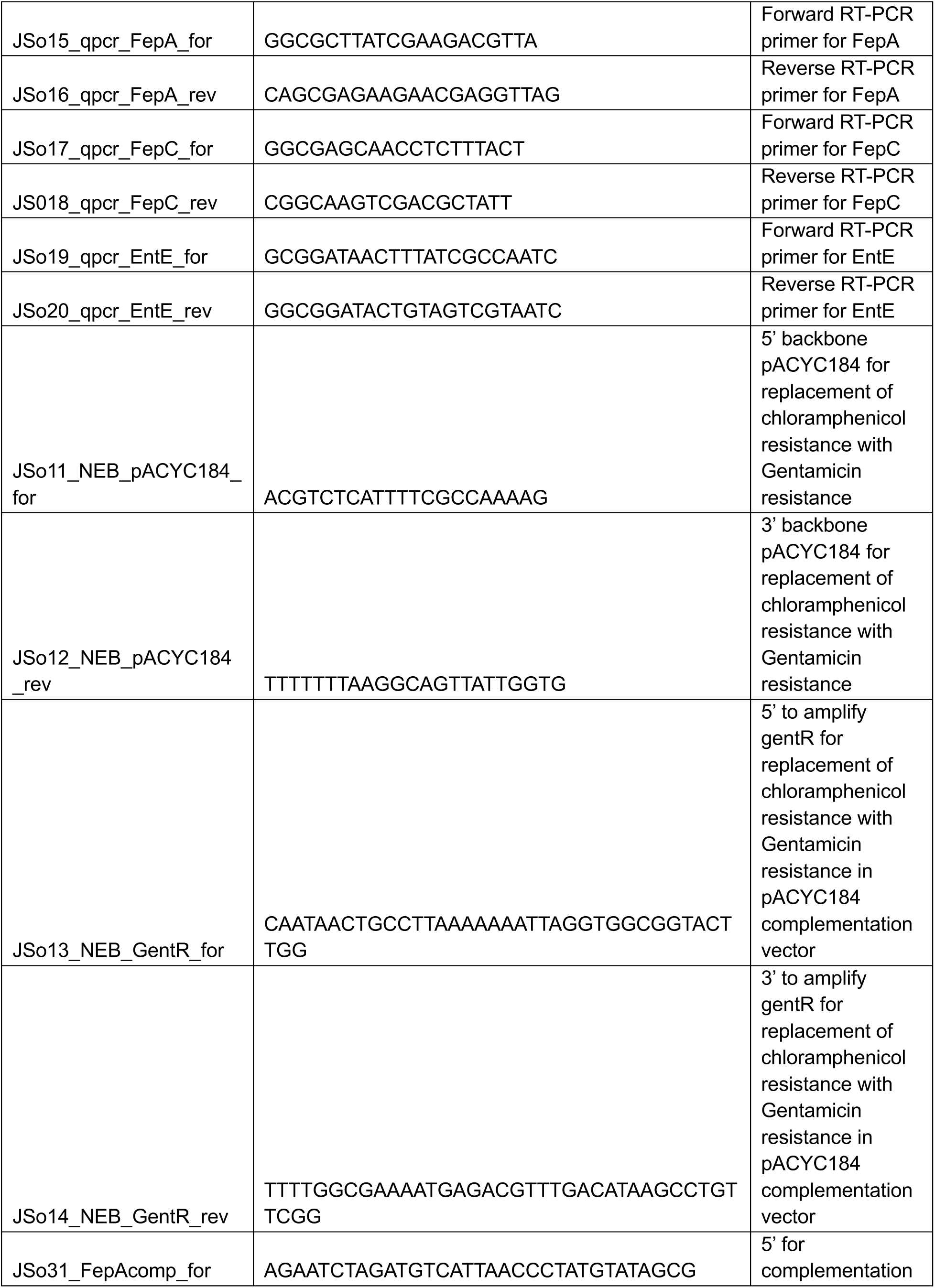
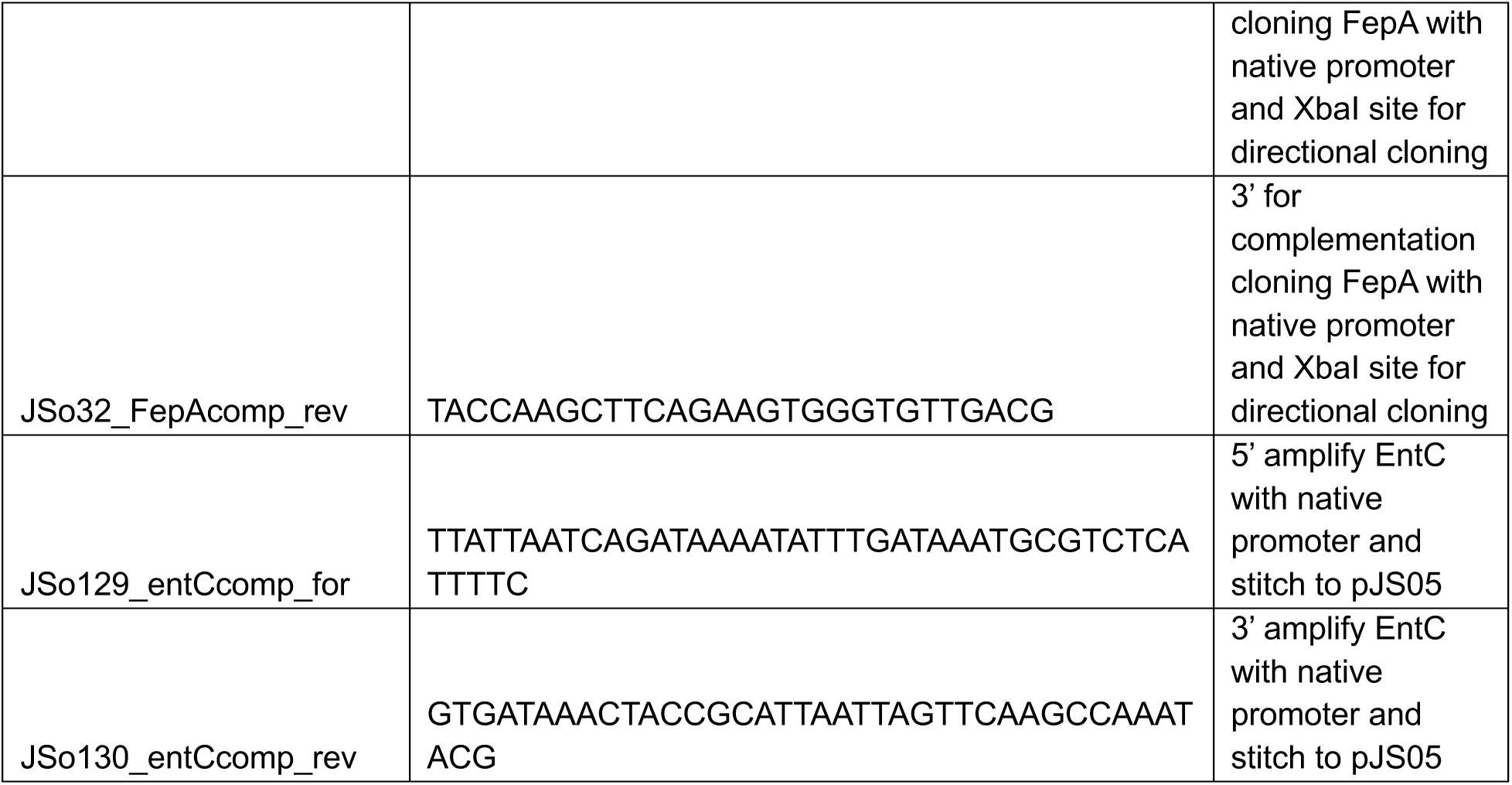
Primers used in this study.

For *fepA* complementation plasmid, empty vector with gentamicin resistance (pJS05) was digested with XbaI and HindIII and genomic DNA from WT KPNIH1 was used as template to amplify *fepA* with native promoter. XbaI and HindIII restriction sites were added to the 5’ and 3’ end of PCR primers respectively for directional cloning using T4 DNA ligase (NEB).

For *entC* complementation vector, genomic DNA from MKP103 was used as template with native promoter for insertion into XbaI and HindIII digested pJS05 vector using HiFi DNA assembly mastermix per manufacturer’s instructions and indicated primers. ∼100ng Empty vector or complementation vectors were transformed into WT and indicated transposon mutant *Kpn* after washing overnight culture 3 times in 300mM Sucrose and electroporating 2.5kV, 25µF capacitance, 200Ω, in 2.0mm gap cuvette with 1h recovery in LB at 37°C before plating on LBA + 10µg/mL gentamicin.

### Arnow’s assay

Assay adapted from Arnow et al. (37, 38). 75µl of supernatant from overnight cultures of WT *Kpn* or transposon mutants grown in Chelexed M9 media was mixed with 75µl of 0.5N HCl in a flat bottom 96 well plate. 75µl nitrite-molybdate reagent (1g NaNO_3_ (Sigma), 1g Na_2_MoO_4_ (Sigma) in 10mL H_2_O) was added and mixed thoroughly before 75µL 1M NaOH added and let incubate for 10 minutes before measuring absorbance at 510nm. Standard curve for “catechol” concentration generated with serial dilutions of 2,3-Dihydroxybenzoic acid (Sigma) and each sample concentration was normalized to overnight OD_600_ measurement before centrifugation to remove supernatant.

### Inductively-coupled plasma mass spectrometry

Overnight cultures were sub-cultured at 1:1000 dilution in 50mL metal-free conical tubes and grown for 3.5h to mid-log growth. Equal volume of EtOH, 50µM TPEN, or 350µM 2-2’ Dipyridyl (Sigma) were added to each culture and returned to grow at 37°C for 1 hour. 1mL culture was washed 2x with PBS in 15mL metal-free conical tube before digestion with 70% Optima-grade nitric acid overnight at 65°C then diluted with UltraPure water to 20% nitric acid for analysis. Elemental quantification was conducted using an Agilent 7700 ICP-MS attached to an ASX-560 autosampler. The settings for analysis were cell entrance = −40 V, cell exit = −60 V, plate bias = −60 V, OctP bias = −18 V, and helium flow = 4.5 ml/min. Optimal voltages for extract 2, omega bias, omega lens, OctP RF, and deflect were empirically determined. Calibration curves for elements were generated using ARISTAR ICP standard mix. Samples were introduced by peristaltic pump with 0.5-mm-internal-diameter tubing through a MicroMist borosilicate glass nebulizer. They were initially taken up at 0.5 rps for 30 seconds, followed by 30 seconds at 0.1 rps to stabilize the signal. Spectrum mode analysis was performed at 0.1 rps, collecting three points across each peak and conducting three replicates of 100 sweeps for each element. The sampling probe and tubing were rinsed with 2% nitric acid for 30 seconds at 0.5 rps between each sample. Data were acquired and analyzed using Agilent MassHunter workstation software version A.01.02.

### qPCR

3 biological replicate overnight cultures were sub-cultured at 1:1000 dilution in 50mL conical tubes and grown for 3.5h to mid-log growth. Cultures were centrifuged at 10min at 4000g and resuspended in 1000µg/mL final concentration CP^ΔS1/S2^ or CP^H3N^ mutant in 40% CP Buffer and 60% LB. Cultures were let grow for 1 hour before 2mL culture was collected and centrifuged 8000g for 2min. Supernatant was removed and cells were resuspended in 1mL Trizol reagent. RNA was extracted as above. 100ng RNA from each sample was used for cDNA synthesis using iScript cDNA synthesis kit (Biorad) per manufacturer’s instructions. cDNA was diluted 1:10 in nuclease free H_2_O and used for 12.5µl qPCR reactions using iQ SYBR Green supermix with primer pairs listed in Table 3. *recA* expression was used as internal control (37) and fold-change values were calculated using ΔΔCT method.

### Enterobactin extraction and high-performance liquid chromatography

WT and *entC::Tn Kpn* strains were grown in 5 ml M9 media overnight. 1mL supernatant, 30µM enterobactin diluted in M9, or equal volume of DMSO diluted in M9 were mixed with 4 mL methanol for one hour at −80°C. Mixture was centrifuged in Beckman Avanti JXN-26 centrifuge at 14000rpm at 4°C for 10 minutes before supernatant was transferred to a new vial and dried under inert gas at 25°C. Samples were reconstituted in 150µl of 5% acetonitrile, heated 5 min at 56°C to dissolve into solution, and ran through a 0.45 uM PTFE filter before analysis on an Agilent 1260 Infinity HPLC. Specifically, 20 ul sample was separated using a Supelco Ascentis Express C18 (50 x 2.1 mm, 5 mm) column with a Phenomenex SecurityGuard (C18 cartridge 3.2 x 8 mm) guard column in place. Mobile phases were made up of 0.1% trifluoroacetic acid in (A) HPLC grade water and (B) HPLC grade acetonitrile. Gradient conditions were as followed: 0 to 5.0 min, B = 5%; 5 to 15 min, B = 50%; 15 to 17 min, B = 100%; 17 to 19 min, B = 0%. The flow rate was maintained at 0.5 mL/min. Absorbance was detected at 220 nm.

### Statistics

Raw data was collected in Microsoft Excel and imported to GraphPad Prism (11.0.2) for statistical analysis and data visualization. Data analyzed as indicated in figure legends and reported N biological replicates for each experiment. Figures were created in Canvas X. Cartoon in Figure 2A Created in BioRender.

## ACKNOWLEDGEMENTS

This work was supported by NIH Grants T32DK007673 to J.R.S., T32HL094296 to D.A.D., T32ES007028 to D.E.K., F31AI197845 to O.S.B., R01AI127793 to W.J.C, and R01AI101171 to E.P.S. and W.J.C. KPNIH1 Arrayed transposon library screen was a generous gift from Dr. Julie Segre. We thank Dr. Michael Grey for WT MKP103 strain. Thank you to the Skaar lab members for critical review of the manuscript and other helpful discussion of the work. We thank Dr. Hayes McDonald at Vanderbilt Mass Spectrometry Resource Center for assistance with proteomics experimental design and implementation. Thank you to Charlotte Singer for assistance with data recording for Arnow’s assay experiments.

## Notes

### Competing Interest Statement

The authors have declared no competing interest.

